# Non-homologous end joining factors XLF, PAXX and DNA-PKcs support neural stem and progenitor cells development

**DOI:** 10.1101/2020.12.01.406629

**Authors:** Raquel Gago-Fuentes, Valentyn Oksenych

## Abstract

Non-homologous end-joining (NHEJ) is a major DNA repair pathway in mammalian cells that recognizes, processes and fixes DNA damages throughout the cell cycle, and is specifically important for homeostasis of post-mitotic neurons and developing lymphocytes. Neuronal apoptosis increases in the mice lacking core NHEJ factors Ku70 and Ku80. Inactivation of other core NHEJ genes, either Xrcc4 or Lig4, leads to massive neuronal apoptosis in the central nervous system (CNS) that correlates with embryonic lethality in mice. Inactivation of one accessory NHEJ gene,e.g. Paxx, Mri and Dna-pkcs, results in normal CNS development due to compensatory effects of Xlf. Combined inactivation of Xlf/Paxx, Xlf/Mri and Xlf/Dna-pkcs, however, results in late embryonic lethality and high levels of apoptosis in CNS. To determine the impact of accessory NHEJ on early stages of neurodevelopment, we isolated neural stem and progenitors cells from mouse embryos and investigated proliferation, self-renewal and differentiation capacity of these cells lacking either Xlf,Paxx, Dna-pkcs, Xlf/Paxx or Xlf/Dna-pkcs. We found that accessory NHEJ factors are important for maintaining the neural stem and progenitor cell populations and neurodevelopment in mammals, which is particularly evident in the double knockout models.

## 1. Introduction

Double-strand DNA breaks (DSBs) are common DNA damage events that threaten the stability of our genome. DSBs can be repaired by homologous recombination (HR), classical non-homologous end-joining (NHEJ) and alternative end-joining (A-EJ, also known as backup end joining, or microhomology-mediated end joining) [1–3]. HR is only available during S/G2 cell cycle phases when the sister chromatid is accessible and then used as a template. C-NHEJ acts throughout the entire cell cycle, sealing directly the broken ends and is the predominant repair pathway in mammalian cells [1, 4]. A-EJ is often microhomology-mediated and more obvious in the absence of classical NHEJ [5].

NHEJ involves recognition of the DSBs by core Ku70/Ku80 heterodimer (Ku), which in turn recruits DNA-dependent protein kinase catalytic subunit (DNA-PKcs) to form a DNA-PK holoenzyme complex that protects free DNA ends. Assembly of DNA-PK triggers the autophosphorylation of DNA-PKcs, as well as DNA-PKcs-dependent phosphorylation of multiple other DNA repair factors [1]. Ku facilitates recruitment of accessory NHEJ factors, such as X-ray repair cross-complementing factor 4 (XRCC4)-like factor (XLF), a paralogue of XRCC4 and XLF (PAXX), and a modulator of retrovirus infection (MRI). Ligation of the broken ends is performed by the core NHEJ factor DNA Ligase 4 (Lig4), which is stabilized by another core factor, XRCC4 [1–3].

Genetic inactivation of *Xrcc4* [6] or *Lig4* [7] in mice results in p53-dependent late embryonic lethality, which correlates with a massive apoptosis in the central nervous system (CNS) [8, 9]. Although *Ku70*^−/−^ and *Ku80*^−/−^ knockout mice are viable, they present high levels of apoptosis in CNS and remarkable growth retardation [10, 11].

In mice, DNA-PKcs, PAXX and MRI are accessory NHEJ factors, while XLF can be considered as either core or accessory factor [2, 3]. *Dna-pkcs*^−/−^ [12], *Xlf^−/−^* [13, 14], *Paxx^−/−^* [15–19] and *Mri*^−/−^ [20, 21] knockout mice are viable, displaying normal growth, lifespan, and neuronal development. However, inactivation of DNA-PKcs kinase domain (*Dna-pkcs*^KD/KD^) leads to Ku- and p53-dependent embryonic lethality, which correlates with high levels of apoptosis in the CNS [22].

More recently, genetic interaction studies uncovered the importance of the accessory NHEJ factors in the development of immune and nervous systems and mouse development in general. Synthetic lethality was reported between *Xlf* and *Dna-pkcs* [19, 23, 24], then between *Xlf* and *Paxx* [2, 15, 16, 18, 19], and finally between *Xlf* and *Mri* [2, 20]. These studies confirmed that functions of DNA-PKcs, PAXX, and MRI are partially compensated by XLF. However, the role of accessory NHEJ factors in early neurodevelopment remains unknown.

Here, using single and double knockout mouse models, we found that XLF, DNA-PKcs and PAXX are required to maintain pluripotency of neural stem cells, including aspects of self-renewal, proliferation, and differentiation to neurons and astrocytes.

## 2. Materials and Methods

### 2.1. Mice

All experimental procedures involving mice were performed according to the protocols approved by the Comparative Medicine Core Facility at Norwegian University of Science and Technology (NTNU, Norway). *Dna-pkcs*^+/−^ [12], *Xlf*^+/−^ [13], and *Paxx*^+/−^ [17] mouse models were previously described.

### 2.2. Mouse genotyping

A conventional polymerase chain reaction (PCR) was used to determine the mouse genotypes. DNA was isolated from ear punches digested overnight at 56°C with 2 % proteinase K in DNA lysis solution containing 10 mM pH=9.0 Tris, 1 M KCl, 0.4 % NP-40 and 0.1 % Tween 20. Next, the samples were heat-treated for 30 minutes at 95°C. The PCR reactions were performed using *GoTaq®G2 Green Master Mix* (Promega, WI, USA; #M7823) or Taq 2x Master Mix Kit (New England Biolabs® Inc., Ipswich, MA, USA; #M0270L) according to the manufacturer’s instructions. Each reaction contained 50 ng of DNA and 0.8 μM of indicated primers (Supplementary Table 5) in a final volume of 25 μL. The PCR product was revealed in a 0.7 % agarose gel.

### 2.3. Neural stem and progenitor cell cultures

Neural stem and progenitor cells (NSPCs) were cultured as free-floating aggregates, also known as neurospheres [21, 25]. Briefly, murine embryos were collected at embryonic day E15.5, the brains were isolated and the cerebellums were removed. Remaining brain parts were mechanically disrupted in proliferation medium, containing Dulbecco Modified Eagle Medium / Nutrient Mixture F12 (DMEM/F12), supplemented with 1 % penicillin/streptomycin, 2 % B27 without vitamin A (Thermo Fischer Scientific, USA), 10 ng/mL EGF and 20 ng/mL bFGF. The neurospheres were formed and incubated at 37°C, 5 % CO_2_ and 95 % humidity. The neurospheres were dissociated every seventh day using 0.25 % of Trypsin in EDTA, as previously described in [21, 25]. For more details, see also *Supplementary Materials and Methods*. The neurospheres from passages 3 to 10 were used in all the experiments.

### 2.4. Proliferation assay

NSPCs proliferation rates were assayed using *PrestoBlue™ Cell Viability Assay* (Thermo Fisher Scientific, Waltham, MA, USA; A13261) following the manufacturer’s protocol and [26]. Briefly, dissociated single NSPCs were loaded onto a 96-well-suspension plate at 8,000 cells/well in proliferation medium. *PrestoBlue*™ reagent was added at 10 % of the well volume at day 3, and incubated for 2 hours at 37°C, 5 % CO_2_ and 95 % humidity. The proportion of live cells was estimated by measuring fluorescence intensity using FLUOstar Omega system (BMG Labtech, Germany).

### 2.5. Self-renewal capacity assay

We followed the protocol described earlier [21]. Briefly, the capacity of neural stem cells to maintain their multipotency *ex vivo* was assessed by determining the number and two-dimensional size of neurospheres. Dissociated single NSPCs were plated onto 6-well suspension plates containing proliferation medium (day 0). At day 8, images of the entire wells were captured using the EVOS microscope (Invitrogen, USA). The pictures were analyzed using the *ImageJ* software (NIH, USA) to obtain the total number of neurospheres per well and size of spheres (pixels, px).

### 2.6. Differentiation assay

Differentiation was induced in dissociated NSPCs, as described previously [21, 25]. Briefly, 25,000 single NSPCs were cultured onto 48-well plates pre-coated with 30 μg/mL poly-D-lysine and 2 μg/mL laminin, with differentiation medium containing NeuroBasal A medium (Thermo Fischer Scientific, USA) supplemented with 1 % penicillin/streptomycin, 2 % B27, 1 % GlutaMAX and 10 ng/mL bFGF (day 0) (also see *Supplementary Materials and Methods*). On day 5, the differentiated cells were fixed with 4 % paraformaldehyde for 15 minutes at room temperature. Further, an immunostaining was performed using antibodies recognizing either the neuron-specific β-III tubulin (Tuj1) or the glial fibrillary acidic protein (GFAP) proteins, to determine neurons and astrocytes respectively after differentiation [21, 25]. Briefly, the cells were permeablized with 0.1 % Triton X-100 for 30 minutes, washed 3 times with PBS (Oxoid Limited, UK), and blocked with 1:2 dilution of blocking solution containing 10 % BSA (Sigma, USA), 10 % goat serum (Invitrogen, USA) and 0.1 % Triton X-100 (Sigma, USA) for an hour, and washed with PBS. Then, the cells were incubated with the indicated primary antibodies in 10 % blocking solution for one hour at room temperature and washed with PBS. Next, the cells were incubated for one hour with the secondary fluorescent marker-conjugated antibodies at room temperature and counterstained with 1 μg/mL of 4’6-diamidino-2-phenylindole (DAPI, *Molecular Probes, USA*). Images were collected using the *EVOS* microscope. Positively-stained cells were counted using *ImageJ* software and presented as a proportion of total cells normalized to WT control.

### 2.7. Western blot

Western blots were performed using antibodies against XLF, PAXX, DNA-PKcs, and β-actin (*Supplementary Materials and Methods*) [17, 26, 27]. Neurospheres were collected and lysed with RIPA buffer (Sigma, USA) containing cOmplete™ EDTA-free Protease Inhibitor Cocktail (Roche, USA) and 1 mM phenylmethanesulfonyl fluoride (PMSF, Sigma, USA). Protein concentrations were determined by Bradford assay (Biorad, USA). Further, 40 μg of protein from each clone was analyzed by the SDS-PAGE gel. Proteins were transferred to the membranes using XCell II™ Blot Module (*ThermoFisher Scientific*, *USA*) at 4°C. Then, the membranes were blocked with 5 % milk in PBS with 10 % Tween 20 (PBST) for one hour at room temperature. Primary antibodies were incubated overnight at 4°C, rinsed with PBST 3 times for 5 minutes and incubated with the secondary antibodies for one hour at room temperature. The blot was washed and incubated with SuperSignal™West Femto (Thermo Fischer Scientific, USA) to reveal the proteins with ChemiDoc™ Touch Imaging System (BioRad, USA).

### 2.8. Statistical analysis

To analyze the data, we pulled together two clones per genotype, representing an independent mouse embryo each, and performed three independent experiments with every clone. All the data shown were normalized to WT average levels. To find statistical differences among the genotypes, Kruskal-Wallis test with Dunn’s multiple comparisons test, as a non-parametric alternative of one-way ANOVA, was used. The statistical analyses were performed using *GraphPad Prism* 7.03 software (GraphPad Prism, USA) [21, 25].

## 3. Results

### 3.1. Impact of XLF, PAXX, and DNA-PKcs on proliferation and self-renewal capacity of neural stem and progenitor cells

Single knockout of NHEJ genes *Xlf*, *Dna-pkcs* or *Paxx* results in viable fertile mice without detectable phenotypes in the CNS [12–18]. Contrary, combined inactivation of *Xlf* and *Dna-pkcs*, or *Xlf* and *Paxx* results in a synthetic lethality (Figure 1A) that correlates with severe apoptosis in the CNS [2, 15, 16, 18, 19, 23, 24]. To further investigate the impact of XLF, DNA-PKcs, and PAXX on the nervous system development, we enriched NSPCs by generating neurosphere cultures from WT, *Xlf*^−/−^, *Paxx*^−/−^, *Dna-pkcs*^−/−^, *Xlf*^−/−^*Paxx*^−/−^, and *Xlf*^−/−^*Dna-pkcs*^−/−^ mouse embryos and characterized cell proliferation, self-renewal, and neural differentiation capacity (Figures 1–3).

**Figure 1.**
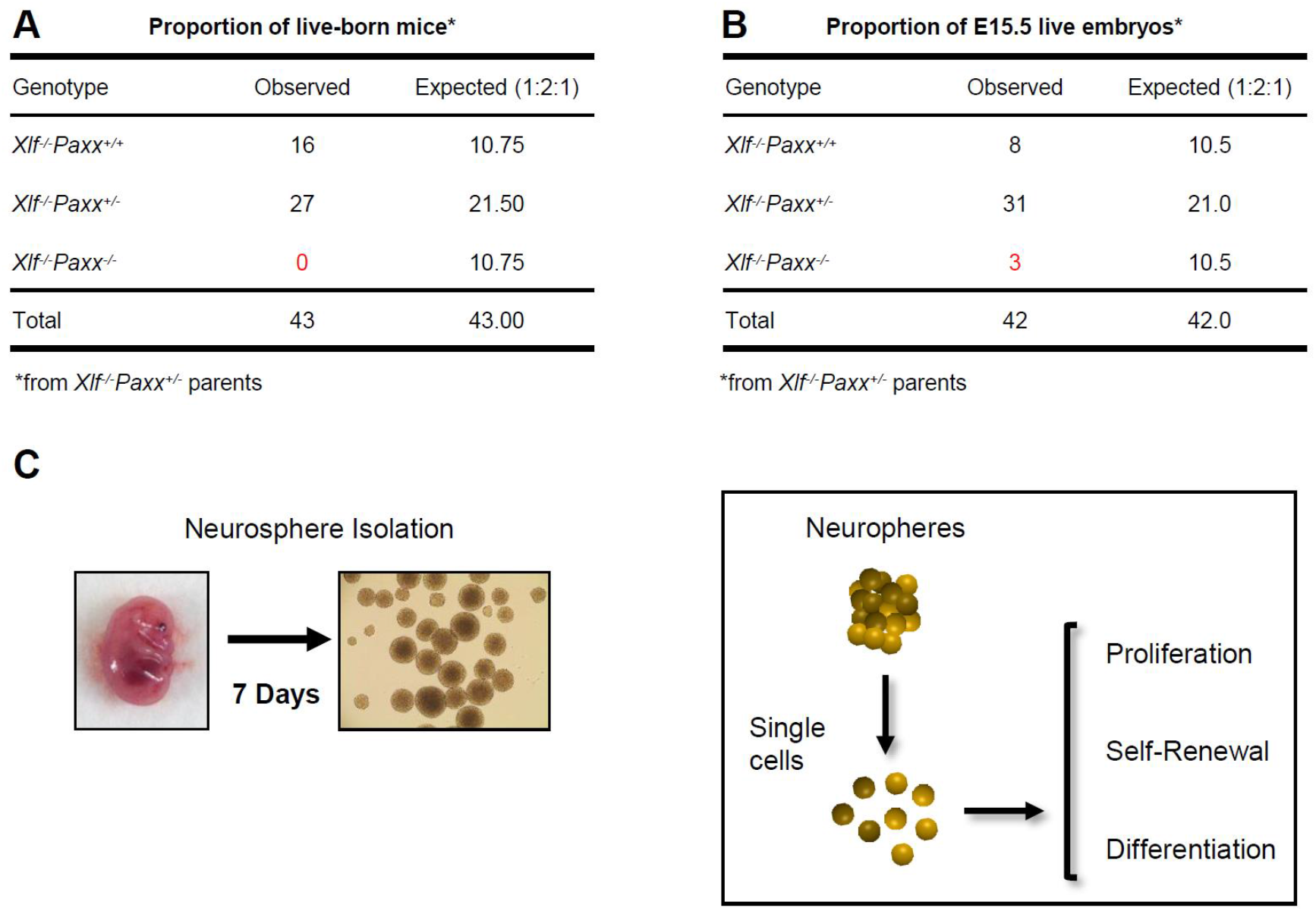
Workflow of the neurosphere-based experiments. **(A)** Synthetic lethality between *Xlf* and *Paxx* in mice. The proportion of live-born mice from *Xlf-*/-*Paxx*^+/−^ parents. No *Xlf*^−/−^*Paxx*^−/−^ double knockout live-born mice were observed out of 43 pups analyzed. (**B)** Fifteen-day-old *Xlf^−/−^Paxx^−/−^* mouse embryos are alive. The proportion of genotypes from *Xlf*−/−*Paxx*^+/−^ parents. Three E15.5 *Xlf*^−/−^*Paxx*^−/−^ embryos were detected out of 42 analyzed. (**C)** Schematic view of the experiment. Embryos were collected at day E15.5 and NSPCs were isolated from the embryonic brains. Single NSPCs formed neurospheres in cell culture. Every seventh days the neurospheres were treated with trypsin to obtain NSPCs used to perform the proliferation, self-renewal and differentiation experiments.

To obtain *Xlf^−/−^Paxx^−/−^* embryos, we intercrossed *Xlf^−/−^Paxx^+/−^* mice. As we observed previously [2, 19], no live-born *Xlf^−/−^Paxx^−/−^* pups were detected (0), while we recorded *Xlf^−/−^Paxx+/+* (16) and *Xlf^−/−^Paxx^+/−^* (27) live-born mice (Figure 1A). By analyzing 15.5E embryos in the same breedings, we detected *Xlf^−/−^Paxx^−/−^* (3), *Xlf^−/−^Paxx+/+* (8) and *Xlf^−/−^Paxx^+/−^* (31) mice, which were later used for the neurosphere generation and characterization. *Xlf*^−/−^*Dna-pkcs*^−/−^ mice were described earlier [19]. Briefly, by breading *Xlf*^−/−^*Dna-pkcs*^+/−^ mice, we obtained no adult *Xlf*^−/−^*Dna-pkcs*^−/−^ mice (0), while there were *Xlf*^−/−^*Dna-pkcs*+/+ (35) and *Xlf*^−/−^*Dna-pkcs*^+/−^ (54) mice at day P30. However, live born *Xlf*^−/−^*Dna-pkcs*^−/−^ mice were detected at days P1-2, in line with our previous observations [19, 23, 24].

By analyzing the neurosphere cultures, we observed that the average proliferation rates of *Xlf*^−/−^ *Paxx*^−/−^ and *Xlf*^−/−^*Dna-pkcs*^−/−^ double knockout neurospheres were reduced when compared to WT and single deficient *Xlf^−/−^*, *Dna-pkcs^−/−^* or *Paxx^−/−^* neurospheres (Figure 2B). To quantify the self-renewal capacity of neurospheres, we plated 10,000 NSPCs and counted the formed neurospheres at day 8 in culture (Figure 2C). Inactivation of *Xlf* resulted in 20% reduction and inactivation of *Paxx* resulted in a 40% reduction of neurosphere count when compared to WT controls. Combined inactivation of *Xlf* and *Paxx* resulted in about 80% reduction of neurosphere count, further highlighting the severe neurological phenotype of *Xlf^−/−^Paxx^−/−^* mice observed *in vivo* [15, 16, 18]. Surprisingly, inactivation of *Dna-pkcs* resulted in a higher number of viable neurospheres, although of smaller size. Combined inactivation of *Xlf* and *Dna-pkcs* resulted in neurosphere count similar to WT controls. We concluded that inactivation of *Xlf* and *Paxx* affected self-renewal capacity and viability of NSPCs (Figure 2C).

**Figure 2.**
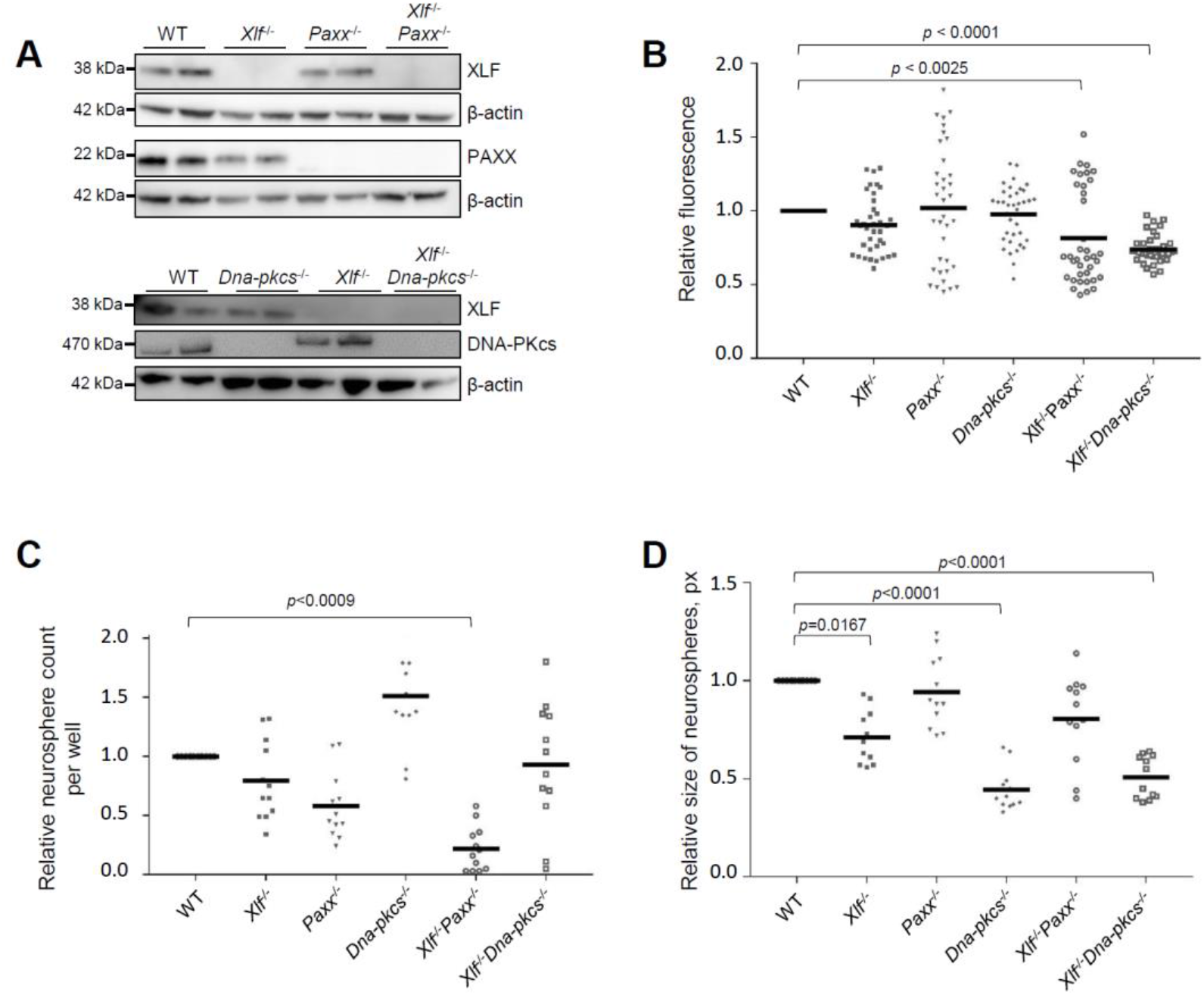
Proliferation and self-renewal capacity of NSPC. (**A**) Western blot analysis revealed no signal corresponding to XLF in *Xlf^−/−^*, *Xlf^−/−^Paxx^−/−^* and *Xlf^−/−^Dna-pkcs^−/−^* NSPC; no signal corresponding to PAXX in *Paxx^−/−^* and *Xlf^−/−^Paxx^−/−^* NSPC; no signal corresponding to DNA-PKcs in *Dna-pkcs^−/−^* and *Xlf^−/−^Dna-pkcs^−/−^* NSPC; β-actin was used as a loading control. (**B**) Amount of neurospheres of indicated genotypes after 48 hours of proliferation, expressed as fluorescence units and normalized to WT controls. Summary of six replicates per two clones, where each clone represents an independent mouse embryo; and three independent experiments (total n=36). The horizontal bars represent the average. Count, n (**C**) and size, pixels (px) (**D**) of neurospheres of indicated genotypes after 8 days in culture. Summary of two replicates per clone, two clones per genotype representing an independent mouse embryos each; three independent experiments (n=12). The horizontal bars represent the average values.

To determine neurosphere growth rate, we used an alternaive quantification based on the image size in pixels (px) (Figure 2D). Inactivation of *Xlf*, *Dna-pkcs,* or both *Xlf/Dna-pkcs*, resulted in neurospheres with 30% to 50% reduction in size when compared to WT controls. Inactivation of *Paxx* did not affect the size of neurospheres in WT and *Xlf*-deficient backgrounds (Figure 2D). We concluded that both XLF and DNA-PKcs support growth of NSPCs in neurospheres.

### 3.2. Impact of XLF, PAXX, and DNA-PKcs on differentiation capacity of neural stem and progenitor cells

To determine whether XLF, PAXX, and DNA-PKcs affect neural differentiation capacity, single NSPCs (25,000 cells) were plated on pre-coated 48-well plates and cultured with differentiation medium for 5 days. Neuronal and glial lineages were identified by immunolabeling using markers for early neurons (Tuj1), and for astrocytes (GFAP). Inactivation of *Xlf, Paxx* or *Dna-pkcs*, and combined inactivation of *Xlf*/*Paxx* did not affect early neuronal differentiation based on average proportions of Tuj1-positive cells (Figure 3A). Combined inactivation of *Xlf* and *Dna-pkcs*, however, resulted in two-fold reduced neurodifferentiation capacity of NSPCs (Figure 3A,C). The proportion of GFAP-positive glial lineage cells increased, although not significantly, when NSPCs were lacking either XLF or PAXX, or both XLF and PAXX (Figure 3B,D).

**Figure 3.**
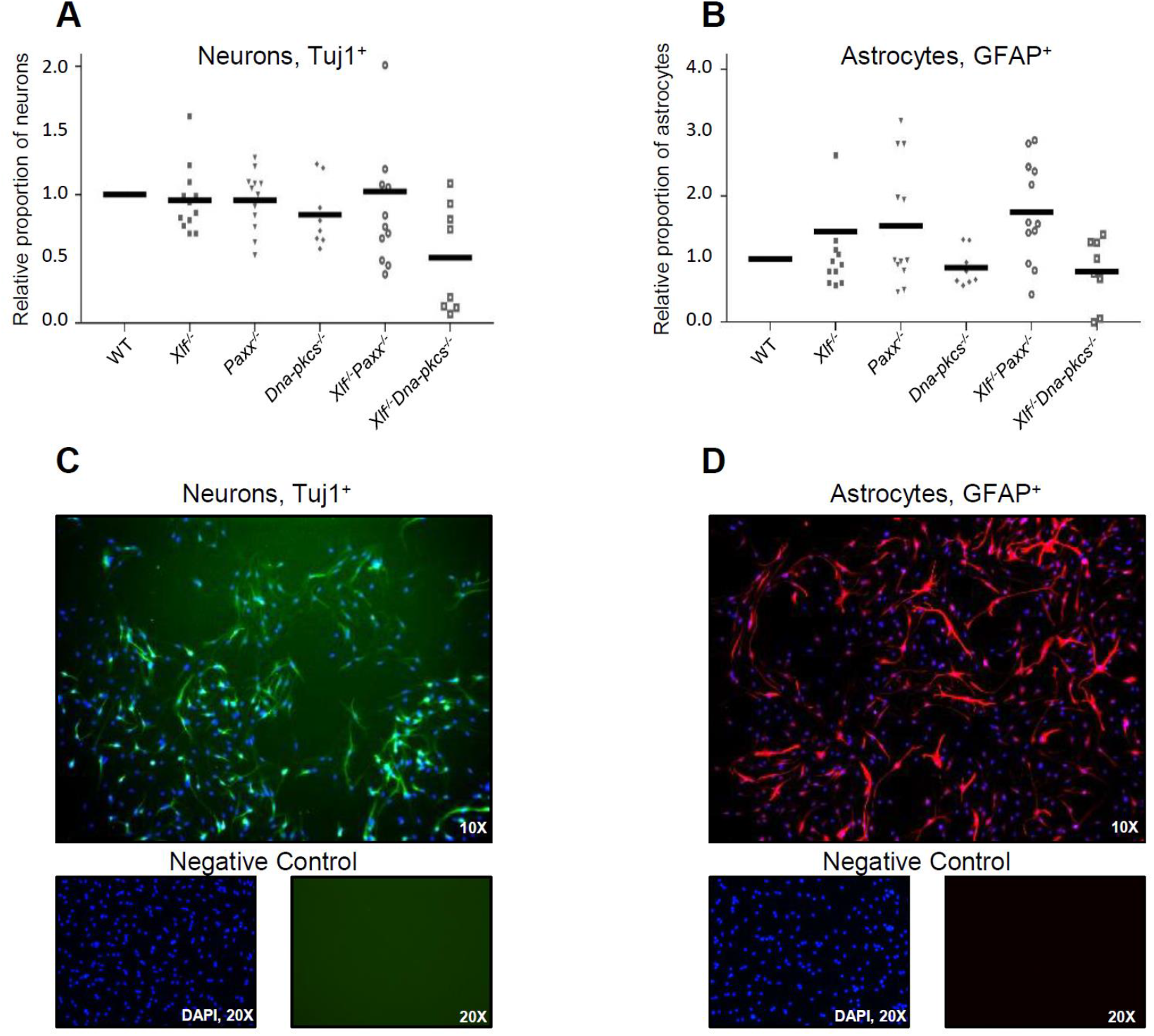
Differentiation of neural progenitors to neurons and astrocytes. (**A**) Proportion of Tuj1^+^ early neurons and (B) GFAP^+^ astrocytes following five days of differentiation from NSPC of indicated genotypes, and normalized by WT controls. Average of two replicates per clone, where two clones represent an independent mouse embryo, and three independent experiments (n=12). **(C, D)** Examples of the immunostaining using antibodies against Tuj1 and GFAP, as indicated. Tuj1^+^ cells are in green (C). GFAP^+^ cells are in red (D). DNA was visualized with DAPI (blue). Negative controls were performed without adding the primary antibodies (bottom).

Overall, XLF possesses functional redundancy with PAXX during the NSPC self-renewal, and with DNA-PKcs during cell growth and neuronal differentiation (Figures 2-3).

## 4. Discussion

Here, we demonstrated that NHEJ factors XLF, PAXX and DNA-PKcs support cellular proliferation during early mammalian neurogenesis, when the proliferation rate is high and the likelihood of DNA damages arising from DNA replication machinery is increased. In *Xrcc4*^−/−^, *Lig4*^−/−^, *Xlf*^−/−^*Paxx*^−/−^ and *Xlf*^−/−^*Dna-pkcs*^−/−^ mice NHEJ is ablated. Therefore, to avoid increased genomic instability during proliferation, developing neurons undergo programmed cell death via the p53-dependent pathway [8, 9, 15, 16, 18, 19, 23, 24].

Although mice lacking XLF possess normal CNS development [13, 14], human patients with mutations in *Cernunnos/XLF* gene suffer from neurological defects, in addition to immunodeficiency [28, 29]. The difference between human and murine phenotypes might be related to the fact that multiple NHEJ and DNA damage response factors, e.g. ATM, H2AX, MDC1, 53BP1, DNA-PKcs, PAXX, MRI, RAG2, partially compensate for the lack of XLF in mice [1, 2, 15, 16, 18–20, 23, 24, 30-37]. In other words, XLF compensates for the lack of multiple factors, including DNA-PKcs and PAXX. Our recent observations revealed that DNA-PKcs and PAXX are likely in the same sub-pathway of NHEJ, because *Dna-pkcs^−/−^Paxx^−/−^* mice do not possess any additional phenotype when compared to the *Dna-pkcs^−/−^* or *Paxx^−/−^* mice [19, 26]. In particular, human HAP1 cell lines lacking both DNA-PKcs/PAXX possess the same levels of genomic instability and sensitivity to DNA damage-inducing agents etoposide, doxorubicine and bleomycin as DNA-PKcs-deficient ones [19, 26]. Moreover, mice lacking both DNA-PKcs and PAXX are live-born, fertile and do not show any additional phenotype when compared to immunodeficient *Dna-pkcs^−/−^* knockout mice [19].

Neurospheres lacking both XLF and DNA-PKcs displayed reduced capacity to differentiate towards neurons that partially explains the severe phenotype of *Xlf^−/−^Dna-pkcs^−/−^* mice [2, 19, 23, 24]. Differently, lack of XLF or PAXX results in a moderate increase in the capacity of neural progenitors to develop into astrocytes. Double knockout *Xlf^−/−^Paxx^−/−^* and *Xlf^−/−^Dna-pkcs^−/−^* neural progenitors possess reduced proliferation capacity (Figure 2B), although due to different reasons. While lack of XLF and PAXX results in lower count of neurospheres likely due to increased rate of cell death (Figure 2C), combined inactivation of *Xlf* and *Dna-pkcs* results in smaller neurospheres (Figure 2D), which can be explained, as one option, by cell cycle arrest due to increased levels of genomic instability [2, 3, 18]. Further analyzes of early neurodevelopment *in vivo* and *in vitro* will help to reveal new insights regarding the role of NHEJ factors in neurodevelopment. Double and multiple-knockout genetic models will facilitate these studies unravelling functional redundancy between the DNA repair factors.

## 5. Conclusions

XLF is functional redundancy with PAXX during the neuronal stem and progenitor cells self-renewal, and with DNA-PKcs during cell growth and neuronal differentiation The NHEJ factors DNA-PKcs, PAXX and XLF are required for an efficient early stage development of neuronal stem and progenitor cells in mice. Additional NHEJ factors such as Mri/Cyren, Ku70, Ku80, XRCC4 and Lig4, as well as multiple ATM-dependent DDR factors might have similar function in neurodevelopment.

## Supplementary Materials

The following are available online at www.mdpi.com/xxx/s1, Table S1: Commercial reagents; Table S2: Antibodies; Table S3: Equipment and Software; Table S4: Solutions and cell culture media; Table S5: Genotyping primers.

## Author Contributions

Conceptualization, R.G.F. and V.O.; methodology, R.G.F. and V.O.; software, R.G.F. and V.O.; validation, R.G.F.; formal analysis, R.G.F.; investigation, R.G.F. and V.O.; resources, V.O.; data curation,R.G.F.; writing—original draft preparation, R.G.F. and V.O.; writing—review and editing, R.G.F. and V.O.; visualization, R.G.F.; supervision, V.O.; project administration, V.O.; funding acquisition, V.O. Both authors have read and agreed to the published version of the manuscript.

## Funding

This research was funded by the following grants: The Research Council of Norway (#249774, #270491 and #291217); the Norwegian Cancer Society (#182355); The Health Authority of Central Norway (#13477 and #38811); The Outstanding Academic Fellow Program at NTNU 2017-2021. VO was a recipient of Karolinska Institutet Stiftelser och Fonder #2020-02155 research grant.

## Acknowledgments

We thank Wei Wang for fruitful discussions during the project development.

## Conflicts of Interest

The authors declare no conflict of interest. The funders had no role in the design of the study; in the collection, analyses, or interpretation of data; in the writing of the manuscript, or in the decision to publish the results.

## Notes

### Competing Interest Statement

The authors have declared no competing interest.

