## Supplemental information for "Non-homologous end joining factors XLF, PAXX and DNA-PKcs support neural stem and progenitor cells development"

**Supplementary Materials and Methods**

### Non-homologous end joining factors XLF, PAXX and DNA-PKcs are required to maintain the neural stem and progenitor cell population

**Raquel Gago-Fuentes[1,2], and Valentyn Oksenych [1,3,4,*]**

[1] Department for Cancer Research and Molecular Medicine (IKOM), Norwegian University of Science and Technology, Laboratory Center, Erling Skjalgssons gate 1, 7491 Trondheim, Norway;

[2] Department of Circulation and Medical Imaging, Norwegian University of Science and Technology, Prinsesse Kristinas gate 3, Akkutten og Hjertelunge-senteret, Postboks 8905, 7491 Trondheim, Norway;

[3] KG Jebsen Centre for B Cell Malignancies, Institute of Clinical Medicine, University of Oslo, N-0316 Oslo, Norway;

[4] Institute of Clinical Medicine, University of Oslo, 0318 Oslo, Norway;

**Key words:** DNA repair; NHEJ; synthetic lethality; genetic interaction

*Corresponding authors:

 (Valentyn Oksenych)

**Supplementary Table S1. Commercial reagents**

| Reagent | Catalogue number, Company, Country |
|---|---|
| *Proteinase K* | #1703001, Invitrogen, Carlsbad, CA, USA |
| *Trizma* | #T3253, Sigma, St. Louis, MO, USA |
| *KCl* | #P9541, Sigma, St. Louis, MO, USA |
| *NP-40* | #127087-87-0, Sigma, St. Louis, MO, USA |
| *Tween-20* | #9005-64-5, Sigma, St. Louis, MO, USA |
| *GoTaq®G2 Green Master Mix* | # M7823, Promega, Madison, WI, USA |
| *DMEM/F12 Medium* | #11330-057,Thermo Fisher Scientific, Waltham, MA, USA |
| *Penicillin/Streptomicyn* | #15140-122, Thermo Fisher Scientific, Waltham, MA, USA |
| *B27 supplement* | #17504044, Thermo Fisher Scientific, Waltham, MA, USA |
| *Epidermal Growth Factor (EFG)* | # AF-100-15, PeproTrech, Sweden |
| *basic-Fibroblast Growth Factor (b-FGF)* | #100-18B, PeproTech, Sweden |
| *Trypsin-EDTA 0.25 %* | #T3924, Sigma, St. Louis, MO, USA |
| *PrestoBlue™ Cell Viability Reagent* | #A13262, Invitrogen, Carlsbad, CA, USA |
| *Neurobasal A Medium* | #10888-022, Thermo Fisher Scientific, Waltham, MA, USA |
| *Poly-D-lysine* | # p0899, Sigma, St. Louis, MO, USA |
| *Laminin* | # L2020, Sigma, St. Louis, MO, USA |
| *B27 supplement without vitamin A* | #12587010, Thermo Fisher Scientific, Waltham, MA, USA |
| *GlutaMAX* | #35050-038, Thermo Fisher Scientific, Waltham, MA, USA |
| *Triton X-100* | #T8787, Sigma, St. Louis, MO, USA |
| *Bovine serum albumin (BSA)* | #A2153, Sigma, St. Louis, MO, USA |
| *Goat antiserum* | #10000C, Invitrogen, Carlsbad, CA, USA |
| *4´6-diamidino-2-phenylindole (DAPI)* | #62248, Molecular Probes, Eugene, OR, USA |
| *RIPA* | #R0278, Sigma, St. Louis, MI, USA |
| *cOmplete™ EDTA-free Protease Inhibitor* | #11873580001, Roche, USA |
| *Phenylmethane sulfonyl fluoride (PMSF)* | #70137720, Sigma, St. Louis, MI, USA |
| *Bradford reagent* | #5000006, BioRad, Hercules, CA, USA |
| *Phosphate-Buffered Saline (PBS)* | #BR0014G, Oxoid Limited, Hampshire, UK |
| *20x NuPAGE Transfer Buffer* | #NP0006-1, Life Technologies, Carlsbad, CA, USA |
| *SuperSignal™ West Femto* | #34095, Thermo Fisher Scientific, Waltham, MA, USA |

**Supplementary Table 2. Antibodies**

| Antibody | Catalogue number, Dilution, Company, Country |
|---|---|
| *Mouse anti-neuron specific β-tubulin (Tuj1)* | #MAB1195, 1:600, R&D Systems, USA |
| *Mouse anti-glial fibrillary acid protein (GFAP)* | #G3893, 1:600, Sigma, USA |
| *Rabbit anti-glial fibrillary acid protein (GFAP)* | #Z0334, 1:1000, Dako, Denmark |
| *Goat anti-mouse Alexa 488* | #A11001, 1:500, Molecular Probes, USA |
| *Goat anti-rabbit Alexa 594* | #A11037, 1:500, Molecular Probes, USA |
| *Rabbit anti-XLF* | #A300-730A , 1:1000, Bethyl, USA |
| *Rabbit anti-C9orf142 (PAXX)* | #126353, 1:200, Novus Biologicals, USA |
| *Mouse anti-DNA-PKCS* | #MA5-13404, 1:1000, Invitrogen, Carlsbad, USA |
| *Mouse anti-β-actin* | #Ab8226, 1:2000, Abcam, UK |
| *Swine anti-rabbit* | #P0399, 1:2000, Dako, Denmark |
| *Goat anti-mouse* | #P0447, 1:2000, Dako. Denmark |

**Supplementary Table 3. Equipment and software**

| Equipment, software | Company, Country |
|---|---|
| *FLUOstar Omega* | BMG Labtech, Ortenberg, Germany |
| *EVOS microscope* | Invitrogen, Carlsbad, USA |
| *ChemiDoc™ Touch Imaging System* | BioRad, Hercules, USA |
| *ImageJ* | National Institute of Health, Bethesda, USA |
| *GradhPad Prism software* | GradhPad Prism, La Jolla, CA, USA |

**Supplementary Table 4. Solutions and cell culture media**

| Solution, medium | Composition |
|---|---|
| DNA lysis solution | 10 mM pH 9 Trizma, 1 M KCl, 0.4% NP-40 and 0.1% Tween20 |
| Proliferation medium | DMEM/F12 medium supplemented with 1% penicillin/streptomycin, 2% B27 without vitamin A, 10 ng/ml EGF and 20 ng/ml bFGF |
| Differentiation medium | NeuroBasal A medium supplemented with 1% penicillin/streptomycin, 2% B27, 1 % GlutaMAX and 10 ng/ml bFGF |
| Blocking solution (10x) | 10% BSA (*Sigma, USA*), 10% goat serum and 0.1% Triton X-100 |
| PBST | 10% Tween20 in PBS |

**Supplementary Table 5. Genotyping primers**

| Gene | Sequence |
|---|---|
| *Xlf* wild type (650 bp) | *Forward: CATGTTGGCTCTGCGAATAGA*<br>*Reverse: GAGCTCGGATATGAGCGCTCAG* |
| *Xlf* knockout (950 bp) | *Forward: CTGTCTTGTGGGCATAGTAGGC*<br>*Reverse: GAGCTCGGATATGAGCGCTCAG* |
| *Paxx* (965 bp wild type; 298, 312, 329 bp knockout) | *Forward: ACAGAGGGTGGTGACTCAGACAATGG*<br>*Reverse: GGAAATGCTATTAGAACCACTGCCACG* |
| *Dna-pkcs* wild type (250 bp) | *Dnapkcs-1: CCCTCCAGACAGCCAGCTAAGACAGG*<br>*Dnapkcs-2: GAAAAAGTCTATGAGCTCCTGGGAG* |
| *Dna-pkcs* knockout (427 bp) | *Dnapkcs-1: CCCTCCAGACAGCCAGCTAAGACAGG*<br>*Dnapkcs-3: ACGTAACTCCTCTTCAGACCT* |
| *Trp53* wild type (321 pb) | *Trp53-1: TGGATGGTGGTATACTCAGAGC*<br>*Trp53-2: AGGCTTAGAGGTGCAAGCTG* |
| *Trp53* knockout (110 bp) | *Trp53-1: TGGATGGTGGTATACTCAGAGC*<br>*Trp53-3: CAGCCTCTGTTCCACATACACT* |
